# Sex Differences in Maturation and Attrition of Adult Neurogenesis in the Hippocampus

**DOI:** 10.1101/726398

**Authors:** Shunya Yagi, Jared E.J. Splinter, Daria Tai, Sarah Wong, Yanhua Wen, Liisa A.M. Galea

## Abstract

Sex differences exist in the regulation of adult neurogenesis in the hippocampus in response to hormones and cognitive training. Here we investigated the trajectory and maturation rate of adult-born neurons in the dentate gyrus (DG) of male and female rats. Sprague-Dawley rats were perfused two hours, 24 hours, one, two or three weeks after BrdU injection, a DNA synthesis marker that labels dividing progenitor cells and their progeny. Adult-born neurons (BrdU/NeuN-ir) matured faster in males compared to females. Males had a greater density of neural stem cells (Sox2-ir) in the dorsal, but not in the ventral, DG and had higher levels of cell proliferation (Ki67-ir) than non-proestrous females. However, males showed a greater reduction in neurogenesis between one and two weeks after mitosis, whereas females showed similar levels of neurogenesis throughout the weeks. The faster maturation and greater attrition of new neurons in males compared to females suggests greater potential for neurogenesis to respond to external stimuli in males and emphasizes the importance of studying sex on adult hippocampal neurogenesis.

**Significance Statement:** Previously studies examining the characteristics of adult-born neurons in the dentate gyrus have used almost exclusively male subjects. Researchers have assumed the two sexes have a similar maturation and attrition of new neurons in the dentate gyrus of adults. However, this study highlights notable sex differences in the attrition, maturation rate and potential of neurogenesis in the adult hippocampus that has significant implications for the field of neuroplasticity. These findings are important in understanding the relevance of sex differences in the regulation of neurogenesis in the hippocampus in response to stimuli or experience and may have consequences for our understanding of diseases that involve neurodegeneration of the hippocampus, particularly those that involve sex differences, such as Alzheimer’s disease and depression.

## 1. Introduction

Adult neurogenesis in the dentate gyrus (DG) has been observed in all mammalian species studied including primates (Kuhn et al., 1996; Gould et al., 1999b; Kornack and Rakic, 1999; Knoth et al., 2010; Briley et al., 2016; Boldrini et al., 2018; Moreno-Jiménez et al., 2019). Despite two papers indicating a lack of neurogenesis in humans (Dennis et al., 2016; Sorrells et al., 2018), recent studies have definitively shown adult neurogenesis exists in humans and is modulated by disease, age, and perhaps sex in response to antidepressants (Epp et al., 2013; Cipriani et al., 2018; Sorrells et al., 2018; Moreno-jiménez et al., 2019; Tobin et al., 2019). Adult hippocampal neurogenesis arises from the radial glia-like neural stem cells (RGLs: type1; Figure 1) in the subgranular zone of the DG, which express stage specific proteins such as Sox2. Sox2 plays a critical role maintaining pluripotency of RGLs (Steiner et al., 2006; Bonaguidi et al., 2011; Encinas et al., 2011; Amador-Arjona et al., 2015; Micheli et al., 2018). The RGLs undergo asymmetrical cell division and generate one RGL and either an astroglia or a transiently amplifying intermediate neural progenitor cell (IPC: type2). The IPCs can undergo multiple symmetrical or asymmetrical cell divisions but generally daughter cells differentiate into neurons (Cameron et al., 1993; Kempermann, 2003; Steiner et al., 2006; Bonaguidi et al., 2011; Encinas et al., 2011). Previous studies show that adult-born cells in the DG divide multiple times, increasing the number of daughter cells which peaks one week after initial mitosis in male rats (Cameron et al., 1993) and perhaps earlier in mice (Amador-Arjona et al., 2015). Adult-born cells in the DG start to die off and show a rapid decrease in the number of new cells between one week and three weeks after the initial cell division in male rodents (Cameron et al., 1993; Snyder et al., 2009; Encinas et al., 2011). A subset of IPCs (type2b), neuroblasts (type3) and immature neurons transiently express a microtubule-associated protein, doublecortin (DCX) for up to three weeks, and new neurons start to express a neuronal nuclear protein, NeuN, approximately one week after mitosis in rats (Brown et al., 2003; Snyder et al., 2009) or two weeks after mitosis in mice (Snyder et al., 2009). Surviving new neurons integrate into the existing neural circuitry, and play an important role in pattern separation and stress resilience (Clelland et al., 2009; Snyder et al., 2011; Hill et al., 2015; França et al., 2017). However, whereas there are species differences in the maturation rate of adult born neurons (Snyder et al., 2009), as of yet no studies to our knowledge have explored sex differences in the maturation rate of adult born neurons.

**Figure 1.**
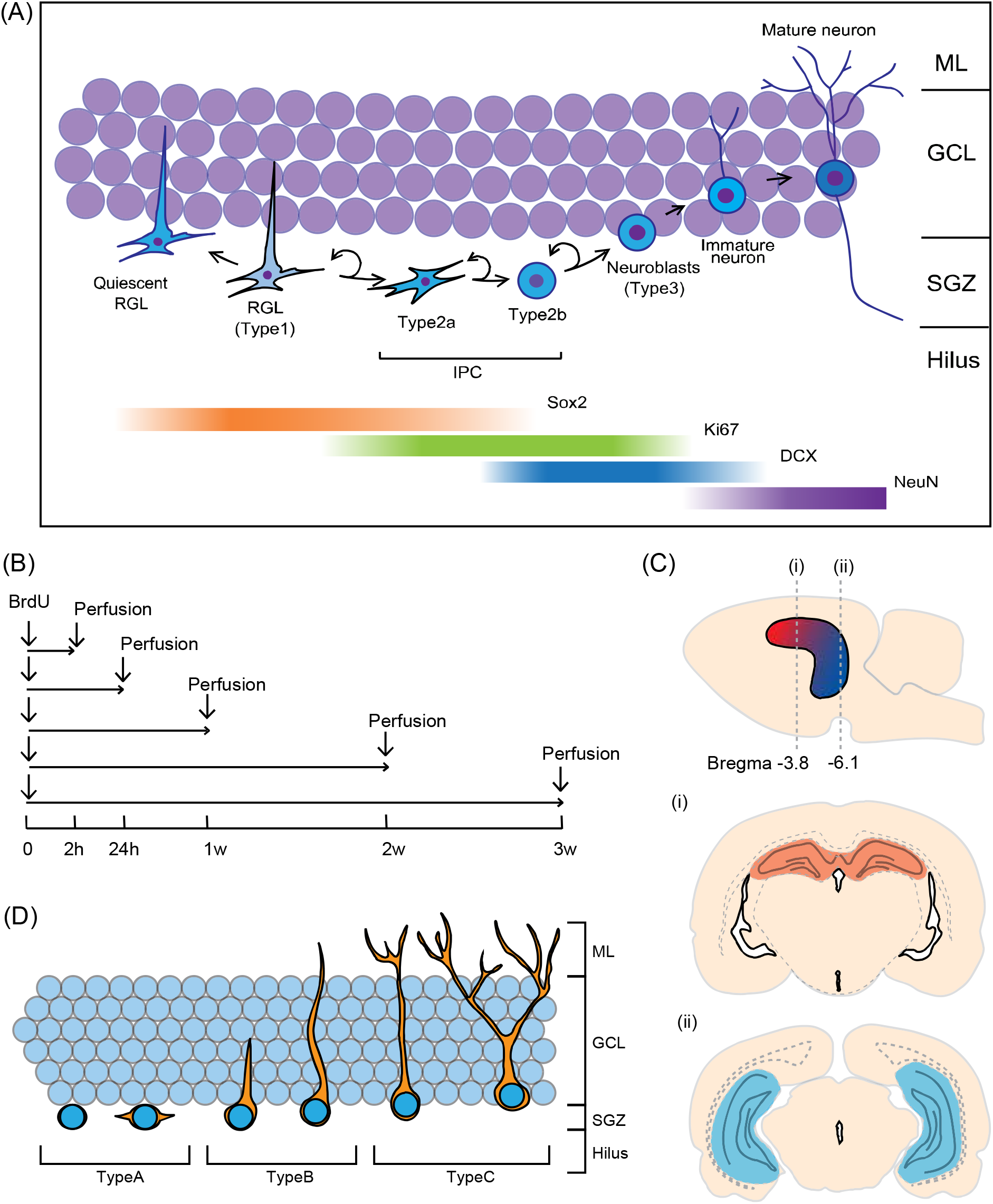
(A) Schematic illustrations for the timeline of neural stem cell lineage with expression of stage-specific proteins (Steiner et al., 2006; Bonaguidi et al., 2011; Encinas et al., 2011; Amador-Arjona et al., 2015; Micheli et al., 2018). (B-D) Schematic illustrations for the experimental design: (B) The experimental timeline, All animals were age-matched and received BrdU injection at 10 weeks. (C) examples of the dorsal (section (i): red; Bregma −3.8mm), and ventral (section (ii): blue; Bregma −6.8mm) hippocampus (numbers represent mm from the bregma) and (D) morphological phenotypes of DCX-ir cells. H- hours, w- weeks, BrdU- bromodeoxyuridine, DCX- doublecortin, GCL- granule cell layer, IPC- intermediate proliferating cell, ML- molecular layer, RGL- radial glial cell, SGZ- subgranular zone

It is important to acknowledge that most of our information about the trajectory and timeline of maturation of new neurons comes from data in male rodents (Cameron et al., 1993; Snyder et al., 2009), with one study in female rodents (Brown et al., 2003). Previous studies demonstrate notable sex differences in the regulation of adult neurogenesis in response to stress, estrogens, androgens, or cognitive training in the DG (Falconer and Galea, 2003; Barker and Galea, 2008; Chow et al., 2013; Hillerer et al., 2013; Yagi et al., 2016; Duarte-Guterman et al., 2019). For instance, acute stress suppresses adult neurogenesis in male rats, but not in female rats (Falconer and Galea, 2003; Hillerer et al., 2013). Furthermore, spatial navigation tasks or spatial pattern separation tasks enhance adult neurogenesis in male rats but not in female rats (Chow et al., 2013; Yagi et al., 2016). The enhancing effect of cognitive training on adult neurogenesis in male rats has a critical period, in which cognitive training must occur 6-10 days after cell birth (Epp et al., 2011), which is curiously the same time that 17β-estradiol also increases neurogenesis in the male meadow vole (Ormerod et al., 2004). The sex differences in the ability of cognitive training to enhance neurogenesis in males but not females suggests one of three scenarios: 1) neurogenesis in the hippocampus is not important for cognitive training in females; 2) the neural activity in the hippocampus may not be as active in females; or 3) there are sex differences in the maturation rate of neurogenesis. Either of these scenarios would lead to the inability of cognitive training to boost survival of new neurons in females in response to spatial training. However, evidence suggests neither of the first two scenarios are correct. Adult DG neurogenesis is associated with better performance in females (Chow et al., 2013; Yagi et al., 2016) and females show increased zif268 expression in the CA3 after training compared to males,(Yagi et al., 2016, 2017).Collectively, these findings suggest sex differences following cognitive training may be due to differences in the maturation rate and perhaps trajectory of adult-born neurons in the DG.

Therefore, the present study aimed to elucidate whether there were sex differences in the maturation and attrition of the new neurons as well as the number of neural stem cells in the dorsal versus ventral DG. A single injection of bromodeoxyuridine (BrdU) was used for birth-dating of adult-born new cells in male and female rats, and brains were immunohistochemically stained for BrdU and endogenous cell-stage-specific protein makers such as Sox2, Ki67, doublecortin (DCX) and NeuN. Given the work above, we expected sex differences in the maturation rate of new neurons with males showing a faster maturation rate than females.

## 2. Materials and Methods

### 2.1. Animals

Forty-four age-matched (two-month old) *Sprague-Dawley* rats were bred at the University of British Columbia and used in this study (n=22 per sex). All subjects were same-sex pair-housed in opaque polyurethane bins (48 × 27 × 20 cm) with paper towels, polyvinylchloride tube, cedar bedding, under a 12h light/dark cycle with 7 am lights-on. Food and water were provided *ad libitum*. Females weighed 240-280g and males weighed 315-355g. All animals were handled every day for two minutes for one week prior to the beginning of the experiment. All experiments were carried out in accordance with Canadian Council for Animal Care guidelines and were approved by the animal care committee at the University of British Columbia. All efforts were made to reduce the number of animals used and their suffering during all procedures.

### 2.2. Experimental design

One intraperitoneal (i.p.) injection of BrdU (200mg/kg) was given to all rats between 11am-12 pm. Rats were perfused either two hours (2h), 24 hours (24h), one week (1w), two weeks (2w) or three weeks (3w) after the BrdU injection, but otherwise were left undisturbed except for weekly cage changes (see Figure 1B). On the day of perfusion, rats were administered an overdose of sodium pentobarbital (500mg/kg, i.p.). Blood samples were collected from the chest cavity, and rats were perfused transcardially with 60 ml of 0.9% saline followed by 120 ml of 4% paraformaldehyde (Sigma-Aldrich). Brains were extracted and post-fixed in 4% paraformaldehyde overnight, then transferred to 30% sucrose (Fisher Scientific) solution for cryoprotection and remained in the solution until sectioning. Brains were sliced into 30 μm coronal sections using a Leica SM2000R microtome (Richmond Hill, Ontario, Canada). Sections were collected in series of ten throughout the entire rostral-caudal extent of the hippocampus and stored in anti-freeze solution consisting of ethylene glycol, glycerol and 0.1M PBS at −20°C until immunostaining. Complete series of sections were immunohistochemically stained for BrdU/DCX and BrdU/NeuN to examine sex differences in the maturation timeline of new neurons, for Sox2 to examine the number of neural stem cells, and for Ki67 to examine actively dividing progenitor cells. In addition, the brain sections were double-stained for BrdU/Sox2 to examine changes of Sox2 expression over the three weeks after BrdU injection.

### 2.3. Radioimmunoassay for 17β-estradiol and testosterone

Previous studies reported that 17β-estradiol increases cell proliferation in females but not males (Tanapat et al., 1999; Barker and Galea, 2008). Androgens increase survival of new neurons in males but not in females, but do not influence cell proliferation in either sex (Spritzer and Galea, 2007; Duarte-Guterman et al., 2019).Thus, we examined serum levels of 17β-estradiol and testosterone in females and males of the 1w, 2w and 3w groups,respectively. Blood samples were stored at 4°C overnight and centrifuged at 10g for 15 minutes to collect serum. Serum 17β-estradiol levels in female rats and serum testosterone levels in male rats were assayed using commercially available radioimmunoassay (RIA) kits from Beckman Coulter (Brea, USA) or MP Biomedicals (Santa Ana, USA) respectively. The sensitivity of the RIA kits was 0.75 ng/mL for 17β-estradiol and 0.03ng/mL for testosterone. The intra- and inter-assay coefficient of variation were <8.9% and <12.2% respectively for 17β-estradiol and <8.2% and <13.2% for testosterone. For females with 50 pg/ml or higher serum estradiol levels were considered to be in proestrus (Cameron et al., 2008). Based on estradiol levels, none of the females in the 1w, 2w and 3w groups were in proestrus at the time of sacrifice (see Table 1).

**Table 1.**
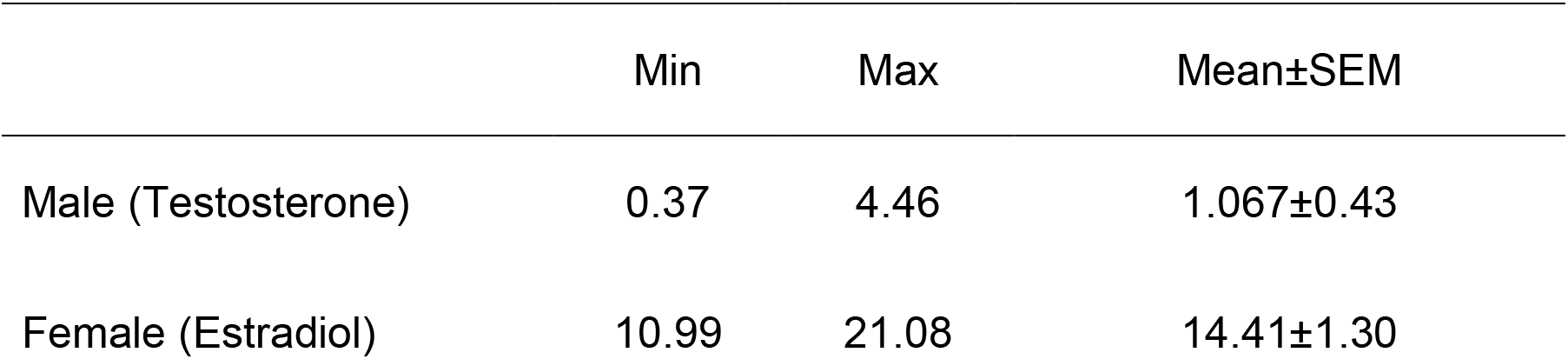
Mean (±SEM), minimum and maximum concentration of serum testosterone in males (ng/ml) and estradiol in females (pg/ml). SEM – standard error of the mean n=13 per group

### 2.4. Estrous cycle stage determination

As the estrous cycle phase can influence cell proliferation (Tanapat et al., 1999; Rummel et al., 2010), estrous cycle stages of the 2h and 24h groups were determined with vaginal lavage samples. Vaginal cells suspended in water were obtained using a glass pipette, transferred onto a microscope slide and stained with cresyl violet (Sigma-Aldrich). Proestrus was determined when 70% of the cells were nucleated epithelial cells. Two females (one each in the 2h and 24h groups) were in proestrus at the time of sacrifice.

### 2.5. Immunohistochemistry

#### 2.5.1. BrdU/NeuN, BrdU/DCX or BrdU/Sox2 double-staining

The exogenous DNA synthesis marker, 5-bromo-2’-deoxyuridine (BrdU) is incorporated into DNA during the synthesis phase of the cell cycle (Kee et al., 2002; Miller et al., 2018). BrdU is a thymidine analogue which is active for two hours after injection in rats (Cameron and Mckay, 2001). Briefly our protocol was as follows: sections were prewashed three times with 0.1 M TBS and left overnight at 4 °C. Sections were then incubated in a primary antibody solution containing 1:250 mouse anti-NeuN (Millipore; MA, USA), 1:200 goat anti-DCX(Santa Cruz Biotechnology; Dallas, Texas, USA) or 1:500 mouse anti-Sox2 (Santa Cruz Biotechnnology; Dallas, Texas USA), 0.3% Triton-X, and 3% normal donkey serum (NDS; Vector Laboratories) in 0.1 M TBS for 24 hours at 4 °C. Next, sections were incubated in a secondary antibody solution containing 1:250 donkey anti-mouse ALEXA 488 (Invitrogen, Burlington, ON, Canada) or donkey anti-goat ALEXA 488 (Invitrogen, Burlington, ON, Canada) in 0.1 M TBS, for 18 hours at 4 °C. After being rinsed three times with TBS, sections were washed with 4% paraformaldehyde for 10 minutes, and rinsed twice in 0.9% NaCl for 10 minutes, followed by incubation in 2N HCl (Fisher Scientific, Waltham, Massachusetts, USA) for 30 minutes at 37 °C. Sections were then rinsed three times in TBS for 10 minutes each and incubated in a BrdU primary antibody solution consisting of 1:500 rat anti-BrdU (AbD Serotec; Raleigh, NC, USA), 3% NDS, and 0.3% Triton-X in 0.1 M TBS for 24 hours at 4 °C. A further incubation of sections commenced in a secondary antibody solution containing 1:500 donkey anti-rat ALEXA 594 (Invitrogen, Burlington, ON, Canada) in 0.1 M TBS for 24 hours at 4 °C. Following three final rinses with TBS, the sections were mounted onto microscope slides and cover-slipped with PVA DABCO.

#### 2.5.2. Ki67 or Sox2 immunofluorescent staining

Ki67 is expressed in actively dividing cells (all stages of the cell cycle except G_0_) and therefore is expressed at higher levels than BrdU 24 h after injection (Kee et al., 2002). Randomly selected brain sections were also immunohistochemically stained with anti-Ki67 or anti-Sox2 (n=8 per sex). Brain sections were prewashed with 0.1 M PBS and left to sit overnight at 4 °C. The next day, sections were incubated in 10mM sodium citrate buffer for 45 minutes at 90 °C to retrieve antigens of Ki67 and blocked with 3% NDS and 0.3% Triton-X in 0.1M PBS, followed by incubation in primary antibody solution made with 1:1000 mouse anti-Sox2 (Santa Cruz Biotechnnology; Dallas, Texas USA) or 1:250 mouse anti-Ki67 (Leica Biosystems; Newcastle, UK), 1% NDS, and 0.3% Triton-X in 0.1 M PBS for 24 hours at 4 °C. Then the sections were incubated in secondary antibody solution, consisting of 1:500 Donkey anti-Mouse ALEXA 488 for Sox2 (Invitrogen, Burlington, ON, Canada) and 1:500 Donkey anti-mouse ALEXA 594 for Ki67 (Invitrogen, Burlington, ON, Canada), 1% NDS, and 0.3% Triton-X in 0.1 M PBS, for 18 hours at 4 °C. After three rinses with PBS, sections were incubated in 1:5000 DAPI in PBS for 3 mins and mounted onto slides and cover-slipped with PVA DABCO.

### 2.6. Cell counting

All counting was conducted by an experimenter blind to the group assignment of each animal using an Olympus epifluorescent microscope and confocal microscope. Location of immunoreactive cells was examined in the dorsal or ventral DG using the criterion defined by Banasr et al. (2006) with sections 7.20-4.48mm from the interaural line (Bregma −1.80 to −4.52mm) defined as dorsal and sections 4.48-2.20 mm from the interaural line (Bregma −4.52 to −6.80mm) as ventral (Banasr et al., 2006; see Figure 1C). Cells were counted separately in each region because the different regions are associated with different functions (reviewed in Fanselow and Dong, 2010) and possibly different maturation timelines (Snyder et al., 2012). The dorsal hippocampus is associated with spatial learning and memory, whereas the ventral hippocampus is associated with stress and anxiety (Moser et al., 1993; Kjelstrup et al., 2002).

#### 2.6.1. BrdU and Ki67

Ki67-ir and BrdU-ir cells were counted under a 100x oil immersion objective lens (Figure 3A, 5A). Every 10^th^ section of the granule cell layer (GCL) that includes the subgranular zone on one half of each brain were counted. An estimate of the total number of cells was calculated by multiplying the aggregate by 10 (Snyder et al., 2005; Ngwenya et al., 2015; Workman et al., 2015). Density of BrdU-ir or Ki67-ir cells was calculated by dividing the total estimate of immunoreactive cells in the GCL by volume of the corresponding region. The volume of the DG was calculated using Cavalieri’s principle (Gundersen and Jensen, 1987) by multiplying the summed areas of the DG by thickness of the section (300μm). Area measurements for the DG were obtained using digitized images on the software ImageJ (NIH).

#### 2.6.2. Percentage of BrdU/NeuN, BrdU/DCX and BrdU/Sox2 co-expression

The percentages of BrdU/NeuN and BrdU/DCX-ir cells were obtained by randomly selecting 50 BrdU-labeled cells and calculating the percentage of cells that co-expressed DCX, NeuN or Sox2 (Figure 5A, 6A and 7A; method used by Banasr et al., 2006). The percentage of BrdU/DCX-ir cells was also categorized into the three morphology types using the criteria used by (Plümpe et al., 2006). Briefly, stages were defined as type-A proliferative: neurons with no or short plump processes, type-B intermediate: neurons possess medium-length processes or apical dendrites that reach the molecular layer, and type-C postmitotic: neurons possess apical dendrites with at least one branching into the molecular layer (see Figure1D). The density of BrdU-ir cells was multiplied by the percentage of BrdU-ir cells that expressed DCX or Sox2.

#### 2.6.3. Sox2

Photomicrographs of the DG were obtained with a 20x objective lens of an Olympus confocal microscope (three images from three sections each from the dorsal and ventral DG; Figure 1C and 2A). Immunoreactive cells were counted automatically using a code developed by JEJS from the digitized images using MATLAB (MathWorks; Natick, Massachusetts, USA). The code is available by contacting the author.

**Figure 2.**
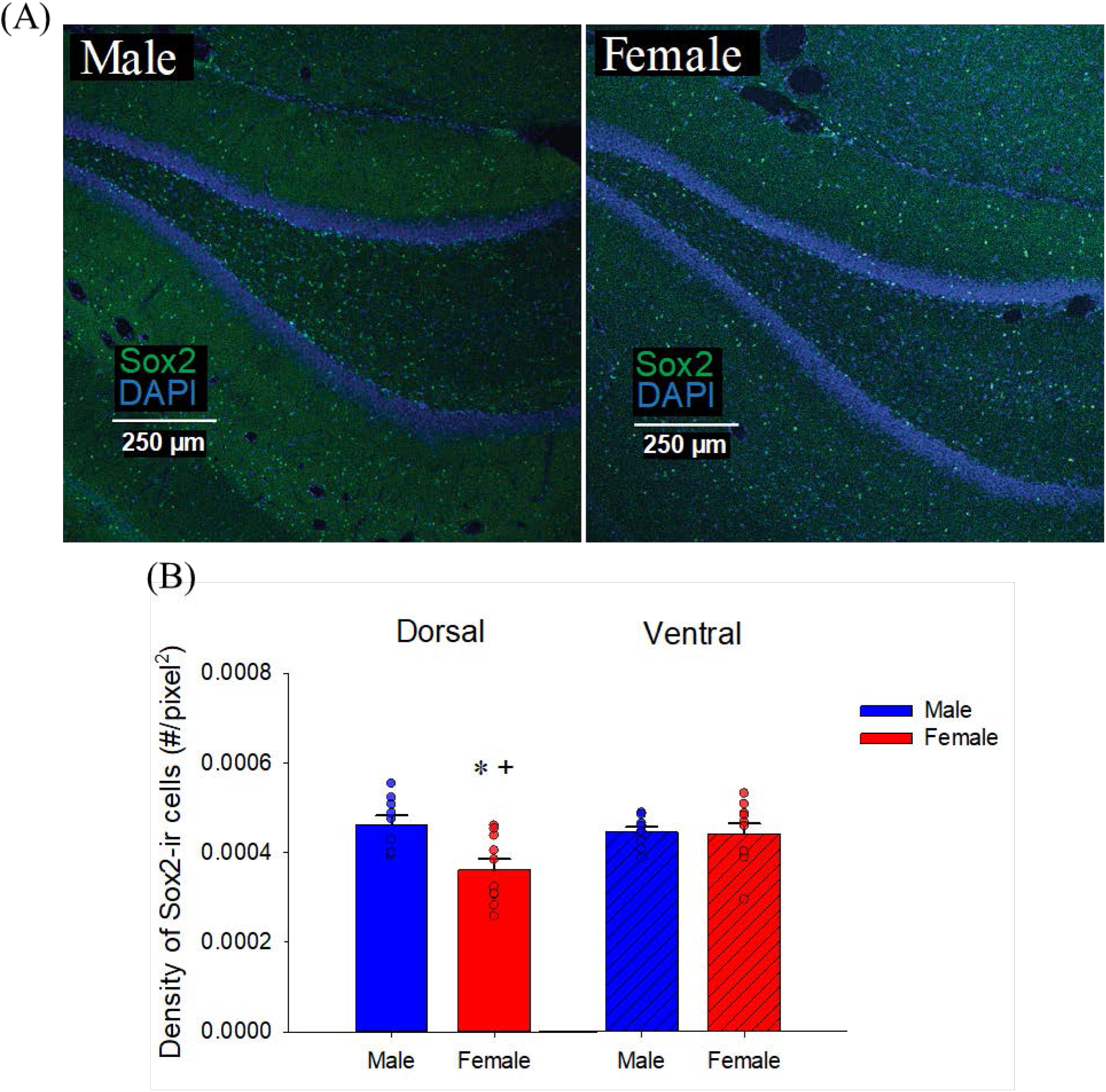
Sex differences in neural stem cells (Sox2-ir). (A) Photomicrographs of Sox2 (green) with DAPI (blue) taken with 10x objective lens from a male (left) and female (right) young adult rat (11 weeks old) in the dorsal dentate gyrus. (B) Mean (+SEM) density of Sox2-ir cells: Males, compared to females, had a greater density of Sox2-ir cells in the dorsal dentate gyrus. The ventral dentate gyrus of females, but not males, had a greater density of Sox2-ir cells compared to the dorsal dentate gyrus. * indicates a significant sex differences and + indicates significant a regional difference (p<0.05). ir- immunoreactive, SEM- standard error of the mean. All animals were age-matched and received BrdU injection at 10 weeks.

### 2.7. Statistical analyses

All analyses were conducted using STATISTICA (Statsoft Tulsa, OK). The density of BrdU-ir cells, BrdU-ir/DCX-ir, or the percentage of BrdU-ir cells that express Sox2 or DCX were each analyzed using repeated-measures analysis of variance (ANOVA), with maturation time (2h, 24h, 1w, 2w, 3w) and sex (male, female) as between-subject variables and with hippocampal region (dorsal, ventral) as the within-subject variable. The percentage of BrdU-ir cells that express NeuN was analyzed using a repeated-measures ANOVA, with maturation time (1w, 2w, 3w) and sex (male, female) as between-subject variables and with hippocampal region (dorsal, ventral) as the within-subject variable. Repeated-measures ANOVAs were used to each analyze the density of Ki67-ir and Sox2-ir cells with sex as between subject factor and with hippocampal region as the within-subject factor. Pearson product–moment correlations were calculated to examine the relationship between dependent variables of interest. Furthermore, the percentage of BrdU/DCX-ir cells expressing type-C morphology was analyzed using repeated-measures ANOVA with sex as between-subject variables and with maturation time and hippocampal region as within-subject variables. Post-hoc tests utilized the Neuman-Keuls procedure. A priori comparisons were subjected to Bonferroni corrections. Significance was set to α=0.05 and effect sizes are given with Cohen’s d or partial η^2^.

## 3. Results

### 3.1. Males had larger dorsal dentate gyrus volumes compared to females

As expected, males had significantly greater volume of dorsal DG compared to females and as such cell density was used for direct comparison between the sexes for all analyses [p = 0.012; region by sex interaction: F(1,22) = 4.61, p = 0.043, Cohen’s d = 1.26; see Table 2]. In addition, the ventral DG was larger than the dorsal DG, as expected [main effect of region: F(1,22) = 36.19, p < 0.0001].

**Table 2.** Mean (±SEM) volume of the dorsal and ventral dentate gyrus in male and female rats (mm^3^). Females had a smaller dorsal dentate gyrus volumes. SEM=standard error of the mean, n= 42 (20 males and 22 females)

|  | Dorsal | Ventral |
| --- | --- | --- |
| Male | 0.905±0.056 | 1.334±0.083 |
| Female | 0.688±0.043 | 1.593±0.195 |

### 3.2. Male rats, compared to female rats, had a greater density of Sox2-ir cells in the dorsal dentate gyrus. Females had greater density of Sox2-ir cells in the ventral compared to dorsal region

To examine sex differences in neural stem cells, we investigated the expression of Sox2. Sox2 is a transcriptional factor that plays a role in maintaining self-renewal of neural stem cells and is considered a neural stem cell marker. Male rats had a greater density of Sox2-ir cells compared to female rats in the dorsal DG (p = 0.024, Cohen’s d = 1.39; sex by region [F(1,16) = 6.34 p = 0.023, see Figure 2B). Females had a greater density of Sox2-ir cells in the ventral DG compared to the dorsal DG (p = 0.005, Cohen’s d = 1.10) whereas this regional difference was not observed in males (p = 0.74). There were trends for a main effect of sex [F(1, 16) = 3.67, p = 0.074] and region [F(1,16) = 4.20, p = 0.057].

### 3.3. Males had greater levels of cell proliferation (Ki67) compared to females

To examine potential sex differences in cell proliferation, we used Ki67, which labels all cells undergoing mitosis. Males had a greater density of Ki67-ir cells compared to females [main effect of sex: F(1,15) = 13.90, p = 0.002, Cohen’s d = 1.80; see Figure 3B]. There was also a trend of main effect of region [F(1, 15) = 3.44, p = 0.083, partial η^2^ = 0.187], but no significant interaction (p=0.11). Because previous studies have observed the rats in proestrus have higher levels of cell proliferation (Tanapat et al., 1999; Rummel et al., 2010), we also examined the relationship between the density of Ki67-ir cells and the levels of 17β-estradiol in females, or testosterone in males, but none was observed (all ps’ > 0.268).

**Figure 3.**
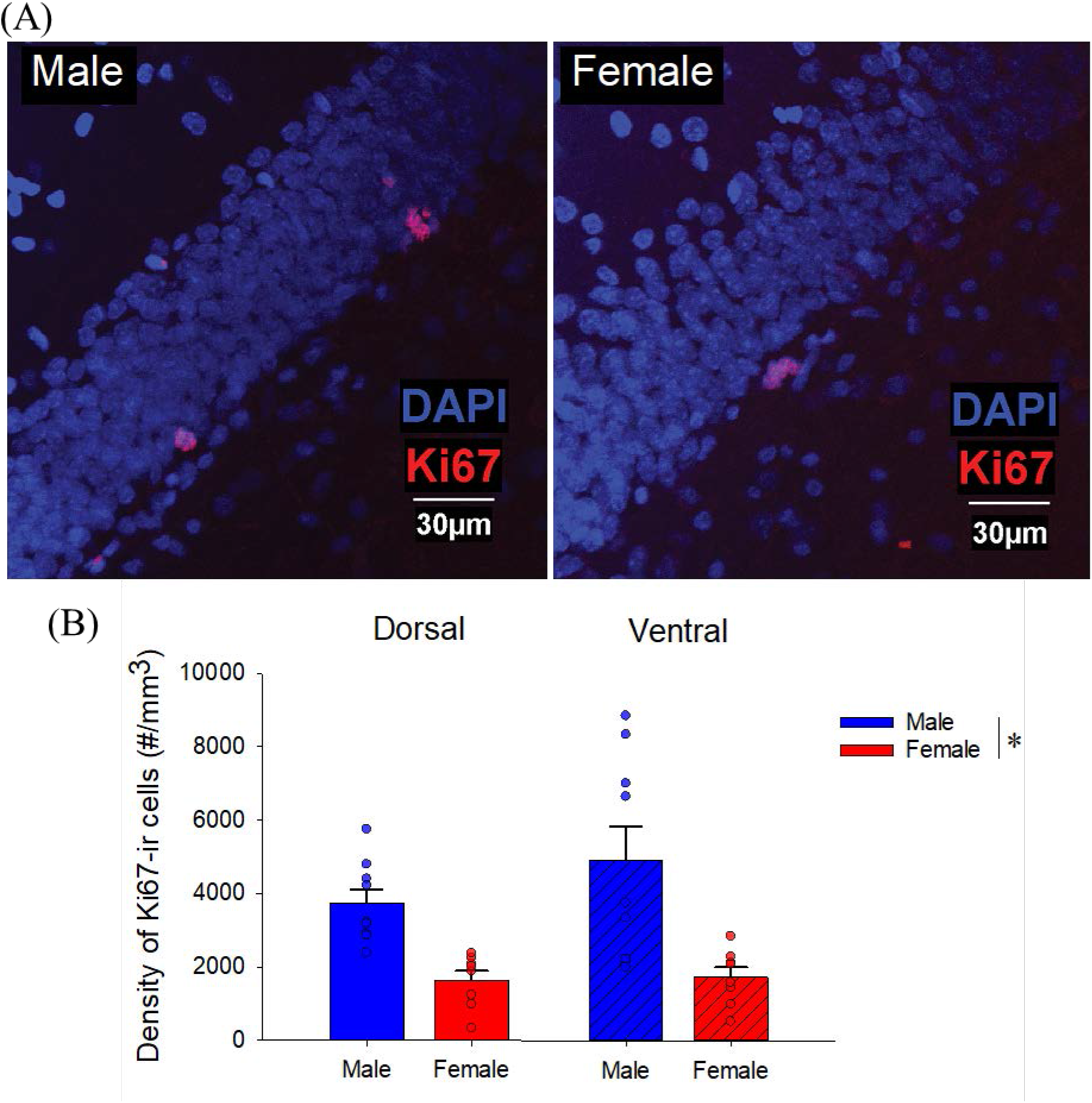
Sex differences in proliferating cells (Ki67-ir) in the dentate gyrus. (A) Photomicrographs of Ki67 (Red) with DAPI (blue) taken with x40 objective from a male (left) and female (right) young adult rat (11 weeks old) in the dorsal dentate gyrus. (B) Mean (+SEM) density of Ki67-ir cells: Males had a greater density of Ki67-ir cells compared to females. * indicates a significant difference (p<0.05). ir- immunoreactive, SEM-standard error of the mean. All animals were age-matched.

### 3.4. Males, but not females, show greater attrition of BrdU-ir cells from 1 week to 2 weeks after mitosis

To determine whether there were sex differences in the trajectory of new neurons across weeks we examined the density of BrdU-ir cells at various time points after BrdU injection (2h, 24h, 1w, 2w, and 3w). Using the same timeline with ^3^H-thymidine, males show an increase ^3^H-thymidine-labelled cells after 24 hours and a large attrition rate of ^3^H-thymidine-labelled from one week to three weeks after injection (Cameron et al., 1993). Consistent with past research (Cameron et al., 1993), males had a greater density of 1w old BrdU-ir cells compared to 2h, 24h, 2w and 3w after BrdU injection (p’s < 0.001; interaction effect of sex by time [F(4,31) = 2.95, p = 0.035, partial η^2^ = 0.276; see Figure 4A]). However, females did not show appreciable differences in the density of BrdU-ir cells across any time points (all p’s > 0.147) except between 2h and 24h (p = 0.156). Furthermore, males had a greater density of BrdU-ir cells than females at the 1w timepoint (p = 0.0003, Cohen’s d = 2.26) but not at any other timepoint (all ps’ > 0.308). Given our findings with Ki67, we also examined sex differences at the 2h and 24h timepoints and saw males had more BrdU-ir cells in the dorsal region only at 2h (priori: p=0.009, Cohen’s d = 2.64) which failed to reach significance at 24 h (p=0.15) compared to females. There were main effects of sex [F(1, 31) = 17.57, p < 0.002, Cohen’s d = 0.746], time [F(4, 31) = 11.78, p < 0.0001, partial η^2^ = 0.603] and region [F(1, 31) = 4.43, p = 0.044, Cohen’s d = 0.254] and an interaction effect of region by time [F(4, 31) = 12.21, p < 0.0001, partial η^2^ = 0.639] was noted but no other significant interactions (p’s > 0.125).

**Figure 4.**
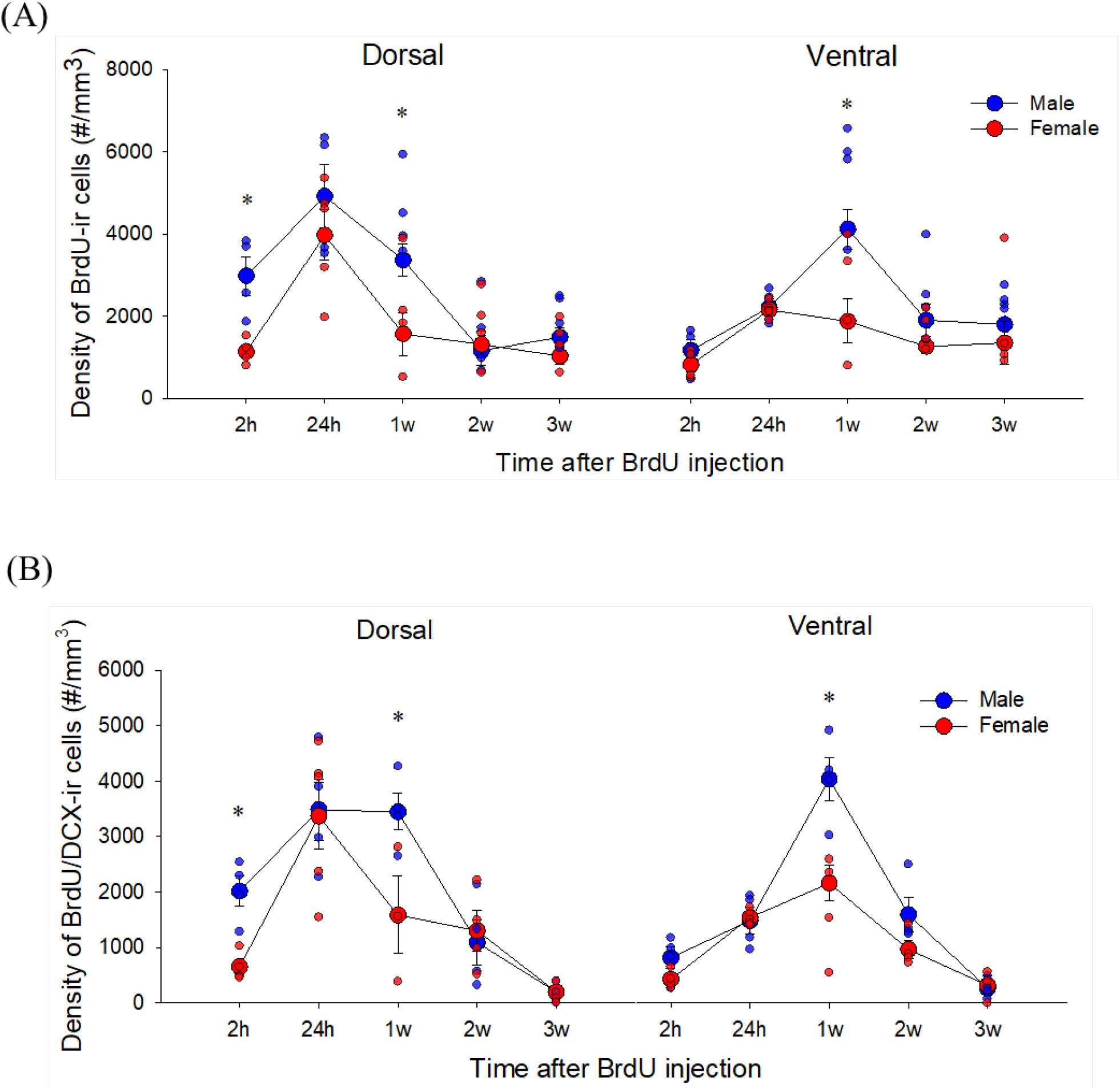
Sex differences in the trajectory of adult-born BrdU-ir cells. (A) Mean (±SEM) density of BrdU-ir cells. Male adult rats had a greater density of BrdU-ir cells at 2h and 1w compared to female adult rats and showed a greater reduction in density between 1w and 2w after BrdU injection. (B) Mean (±SEM) density of BrdU/DCX-ir cells. Males had a greater density of BrdU-ir cells that express DCX cells at 2h and 1w. * indicates a significant sex difference (p<0.05). h-hours, w-weeks, BrdU- bromodeoxyuridine, DCX- doublecortin, SEM-standard error of the mean. All animals were age-matched and received BrdU injection at 10 weeks.

Complementing the attrition rate in BrdU-ir cells across weeks in males, we found that males had a greater density of BrdU/DCX-ir cells than females only at the 1w time point (p=0.00036, Cohen’s d = 2.61) but not at any other timepoint (all p’s > 0.130 [interaction effect of sex by time: F(4, 29) = 4.04, p = 0.0101, partial η^2^ = 0.358; see Figure 4B]. Given our findings with Ki67, we also examined the 2h and 24h timepoint and found that males had a greater density of BrdU/DCX-ir cells compared to females in the dorsal dentate gyrus at 2h (p = 0.005, Cohen’s d = 3.18). There were also main effects of sex [F(1,29) = 11.71, p = 0.0047, Cohen’s d = 0.320], time [F(4, 29) = 29.31, p < 0.0001, partial η^2^ = 0.802] and region [F(1, 29) = 8.66, p = 0.0063, partial η^2^ = 0.230] and an interaction effect of region by time [F(4, 29 = 12.86, p < 0.0001), partial η^2^ = 0.639] but no other significant interactions were noted (p’s > 0.269).

### 3.5. Male adult-born neurons mature faster compared to female adult-born neurons

We then examined whether there are sex differences in maturation rate of adult-born neurons by examining the percentage of BrdU-ir cells expressing maturation stage specific neuronal markers, immature neurons (DCX) and mature neurons (NeuN) across the three weeks. Males, compared to females, had a greater percentage of BrdU-ir cells that expressed NeuN 2w (p = 0.003, Cohen’s d = 2.14) but not 1w (p=0.99) or 3w (p=0.54) after BrdU injection (interaction effect of sex by time [F(2, 17) = 3.52, p = 0.05, partial η^2^ = 0.293; see Figure 5B]). There were also main effects of sex: [F(1, 17) = 7.14, p = 0.016, partial η^2^ = 0.296] and time [F(2, 16) = 41.92, p < 0.00001, partial η^2^ = 0.834] but no other significant main or interaction effects (all p’s > 0.24). The percentage of BrdU-ir cells that expressed NeuN by three weeks after BrdU injection in both males and females was approximately 90% and did not significantly differ between the sexes (p = 0.583).

**Figure 5.**
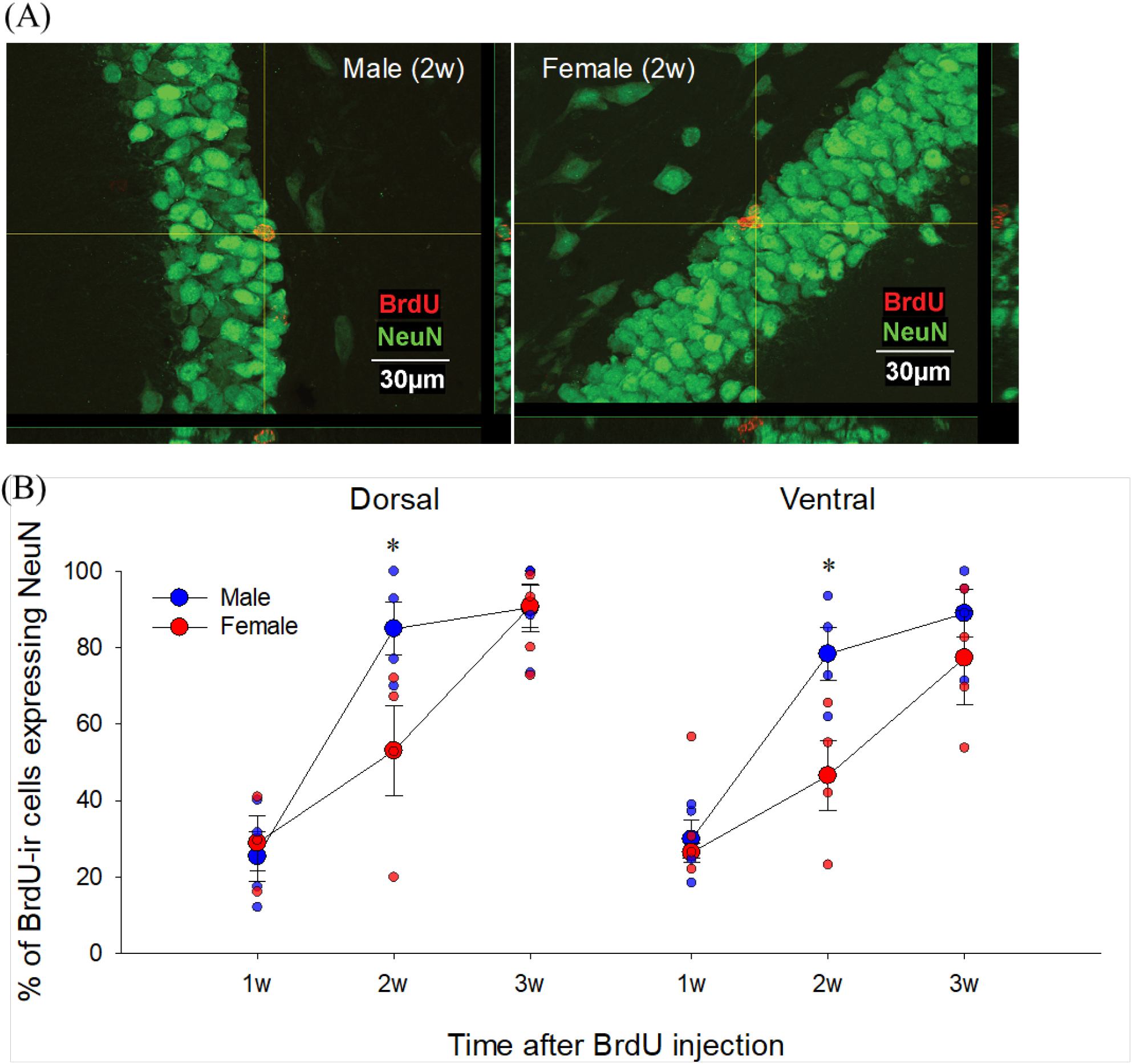
Sex differences in the maturation rate of adult-born neurons in the dentate gyrus (BrdU/NeuN). (A) Photomicrographs of BrdU (red)/NeuN (green) taken with 60x objective lens from a male (left) and female (right) young adult rats in the 2w group. (B) Mean (±SEM) percentages of BrdU-ir cells that express NeuN. Male young adult rats had a greater percentage of BrdU-ir cells that express NeuN at 2w in the dorsal and ventral dentate gyrus. * indicates a significant sex difference (p<0.05). w-weeks, BrdU- bromodoxyuridine, ir- immunoreactive, SEM-standard error of the mean. All animals were age-matchedand received BrdU injection at 10 weeks old.

As expected, in both sexes across both regions, the percentage of BrdU-ir cells that also express DCX decreased significantly as time progressed with the least co-expression at 3w compared to all other time points (all p’s <0.002). Furthermore, the 2h timepoint had lower co-expression than all other earlier timepoints (all p’s <0.024) except 2w (p=0.34) and 3 w [main effect of time: F(4, 30) = 63.69, p < 0.0001; partial η^2^ = 0.895; see Figure 6B]. Females had greater percentage of BrdU-ir cells that co-expressed DCX in 24h group compared to 2h group (a priori: p = 0.0003, Cohen’s d = 6.68; see Figure 6B), which was not seen in males (p = 0.895; sex by time interaction (p = 0.086)). There were no other significant main or interaction effects on the percentage of BrdU-ir cells that co-express DCX (p’s > 0.12). Given the findings showing that new neurons expressed NeuN faster in males compared to females, we also examined BrdU/DCX-ir cells by maturation stage, which we classified using morphology (Plümpe et al., 2006). Consistent with our BrdU/NeuN findings, males had a greater percentage of BrdU/DCX-ir cells expressing type-C morphology compared to females at 2w in the dorsal DG [a priori: p = 0.017, Cohen’s d = 1.84; effect of time: F(2, 18) = 5.39, p = 0.015, partial η^2^ = 0.37; see Figure 6C) but not at 1w (p = 0.95) or 3w (p = 0.84) after BrdU injection.

**Figure 6.**
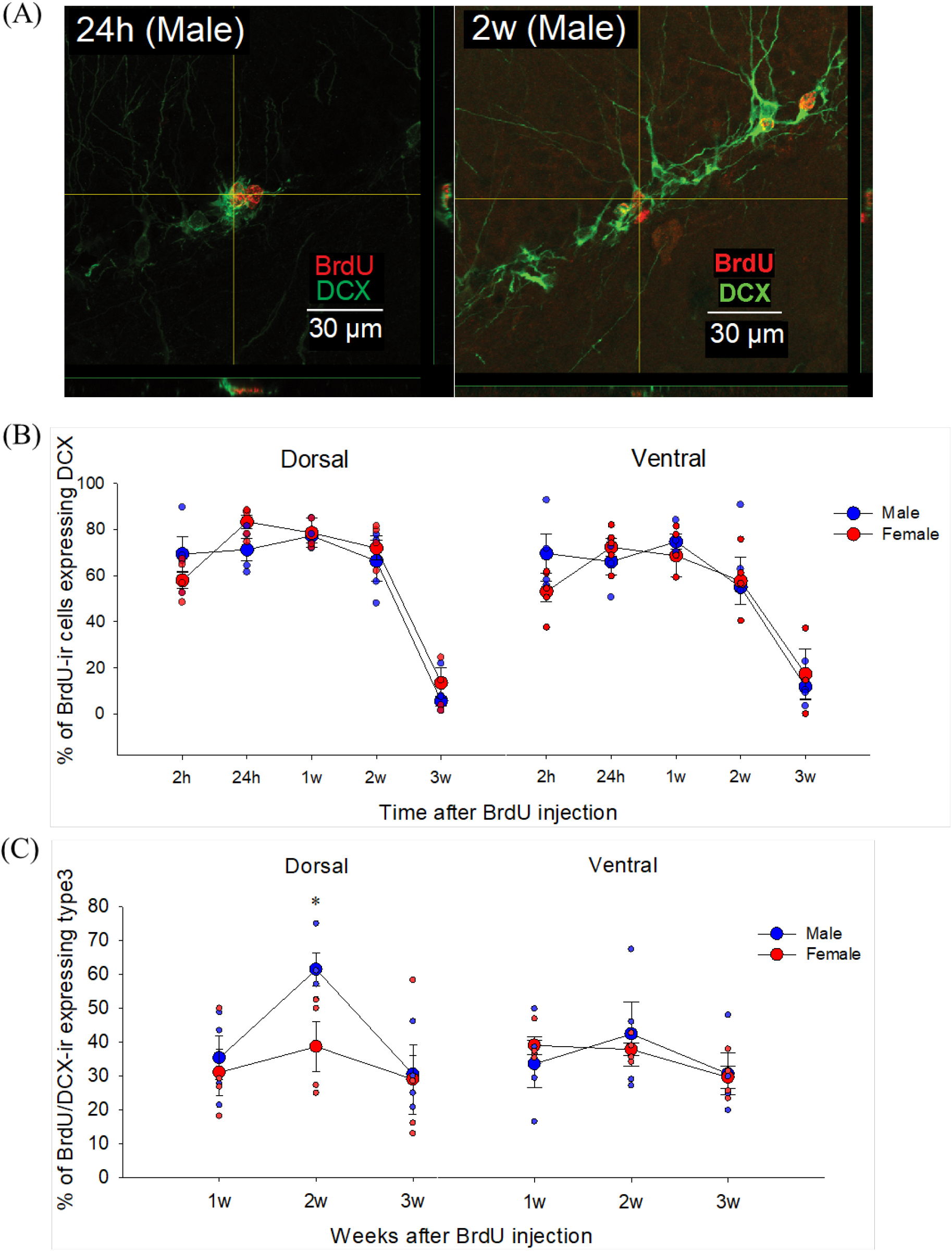
Sex differences in the maturation rate of adult-born neurons in the dentate gyrus (BrdU/DCX). (A) Photomicrographs of BrdU (Red)/DCX (Green) taken from male young adult rat at 24h (left: 60x objective lens) and 2w (right: 40x objective lens) group. (B) Mean (±SEM) percentages of BrdU-ir cells that express DCX. There was no significant sex difference in the percentage of BrdU-ir cells that co-express DCX (C) Mean (±SEM) percentages of BrdU/DCX-ir cells that had a type-C morphological phenotype. A priori comparisons showed that male adult rats had a greater percentage of BrdU/DCX-ir cells that showed the type-C morphological phenotype at 2w compared to female adult rats in the dorsal dentate gyrus. * indicates a significant sex difference (p<0.05). h-hours, w-weeks, BrdU- bromodeoxyuridine, DCX- doublecortin, ir- immunoreactive. All animals were age-matched and received BrdU injection at 10 weeks old.

### 3.6. Males have a greater density of BrdU/Sox2-ir cells in the dorsal DG at 2h compared to females

To understand if there are differences between sexes in the time course of neural stem cell marker expression after mitosis, we examined the density of BrdU/Sox2-ir cells at 2h, 24h, 1w, 2w and 3w after BrdU injection. Males had a greater density of BrdU/Sox2-ir cells compared to females in the dorsal dentate gyrus at 2h but not at any other timepoint [a priori: p = 0.0019; see Figure 7B]. In addition, the dorsal dentate gyrus had a greater density of BrdU/Sox2-ir cells at 2h and 24h than the ventral dentate gyrus compared to all other timepoints (all p’s <0.0003; interaction of region by time F(4, 31) = 11.66, p < 0.0001, partial η^2^ = 0.601). There were also significant main effects of time (F(4, 31) = 40.46, p < 0.0004, partial η^2^ = 0.84) and region (F (1, 31) = 20.50, p < 0.0001, partial η^2^ = 0.398) but no other main or interaction effects (both p’s >0.109).

**Figure 7.**
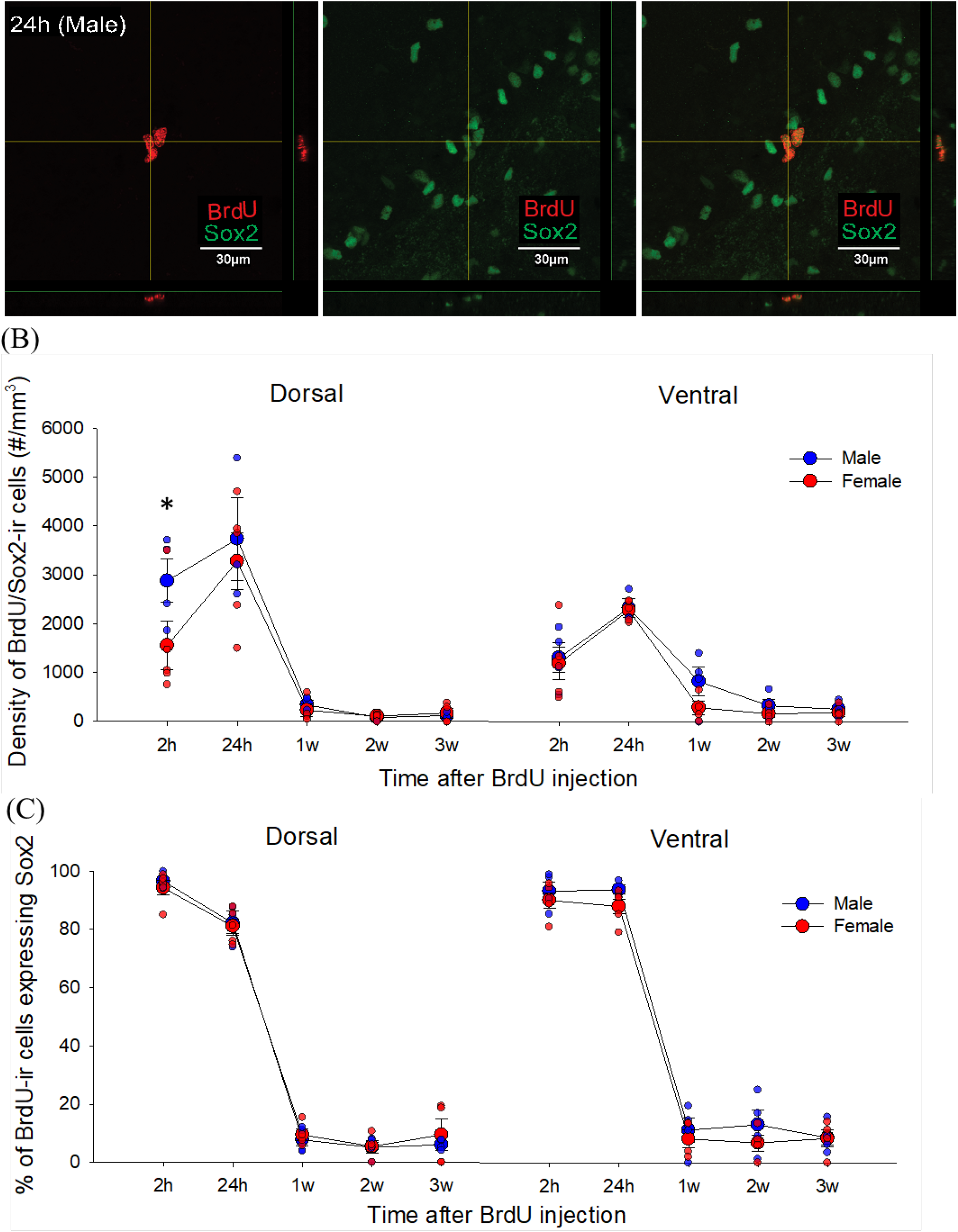
Sex differences in BrdU/Sox2-ir cells across timepoints. (A) Photomicrographs of BrdU (left: red) /Sox2-ir (center: green) cells and merged images (right), taken from a male young adult rat in 24h group. (B) Mean (±SEM) density of BrdU-ir cells that express Sox2. A priori comparisons showed that male, compared to female, young adult rats had a greater density of BrdU-ir cells that co-expressed Sox2 in the dorsal dentate gyrus at 2h after BrdU injection. * indicates a significant sex difference (p<0.05). BrdU- bromodeoxyuridine, ir- immunoreactive, SEM-standard error of the mean. All animals were age-matched and received BrdU injection at 10 weeks old.

### 3.7. The percentage of BrdU/Sox2 co-expressing cells decreased dramatically over time in both sexes

As expected, the percentage of BrdU-ir cells expressing Sox2 decreased across time, with the highest levels at the 2h and 24h timepoints in the dorsal and ventral region (all p’s < 0.0002), with the 2h timepoint having higher levels than 24h in the dorsal dentate gyrus only (p = 0.003; interaction effect of region by time: F(4, 31) = 4.25, p = 0.007, partial η^2^ = 0.354; main effect of region: F(1, 31) = 5.37, p = 0.027, partial η^2^ = 0.148; main effect of time: F(4, 31) = 640.85, p < 0.001, partial η^2^ = 0.988; see Figure 7C]. There was a trend for an interaction effect of region by sex [F(1, 31) = 3.77, p = 0.061, partial η^2^ 0.108]. There were no other significant main or interaction effects on the percentage of BrdU-ir cells expressing Sox2 (p > 0.317).

## 4. Discussion

Our findings indicate that adult-born neurons mature faster in males compared to females. We also found notable sex differences in the attrition or survival rate of BrdU-ir cells across weeks, with males showing reductions across time, and females showing no appreciable reduction in the density of BrdU-ir cells across the three weeks. Furthermore, males had a higher density of dorsal neural stem cells (Sox2) and cell proliferation (Ki67) compared to females. There were notable differences in early expression of DCX in females, but not in males, showing a greater percentage of BrdU-ir cells expressing DCX at 24h compared to 2h. Intriguingly, the density of BrdU-ir cells 2 weeks after production was comparable between males and females. Although a tremendous amount of research has unveiled the characteristics of neurogenesis in the adult hippocampus, these findings underscore that we cannot assume that the same characteristics will be similar in females as they are in males.

### 4.1. Male adult-born dentate granule cells mature faster compared to female adult-born dentate granule cells

We found that adult born neurons mature faster in males than in females, with males showing a rapid increase in the percentage of BrdU-ir cells that expressed NeuN at 2 weeks. Although previous studies did not directly compare the sexes, they are consistent with our results (Brown et al., 2003; Snyder et al., 2009). These studies showed that in male rats 65-75% of BrdU-ir cells expressed NeuN two weeks after BrdU injection (Snyder et al., 2009), whereas a separate study found in female rats less than 10% of BrdU-ir cells expressed NeuN at two weeks after BrdU injection (Brown et al., 2003). Sex differences in the maturation time course of new neurons may be due to sex differences in the neural activity of the hippocampal network. Maturation of adult-born neurons is accelerated by electrophysiological activity in the hippocampus (Piatti et al., 2011), and cFos expression in the dorsal CA3 of hippocampus is greater in males compared to females in response to a Morris water maze task and radial arm maze task (Yagi et al., 2016, 2017). However, in the same studies, females show greater activation of zif268 in the dorsal CA3 compared to males, which is inconsistent with the interpretation of greater activity in the hippocampus accounting for the sex differences in maturation timelines. Another possible explanation for the higher percentage of more mature adult-born neurons in males compared to females at two weeks may involve competition and/or apoptosis resulting in part from the greater attrition from one to two weeks in males, which may impact the survival rate of new neurons (Bergami and Berninger, 2012). Further research is needed to examine the mechanisms of the sex differences in the maturation of new neurons.

### 4.2. Males had more neural stem cells than females, whereas females showed a regional difference with more neural stem cells in the ventral, compared to dorsal, dentate gyrus

In the present study, males had a greater density of Sox2-ir cells in the dorsal DG compared to females. We also found that females had a greater density of Sox2-ir cells in the ventral compared to the dorsal region, that was not observed in males. To our knowledge, neither of these findings have been reported previously. These findings suggest that within females, there is more chance of maintaining pluripotency in the ventral compared to the dorsal DG. How this might be reflected in sex differences in the functions attributed to the dorsal versus ventral hippocampus remains to be determined. However, there are some intriguing possibilities as males generally show better spatial learning (Jonasson, 2005; Voyer et al., 2017), whereas females show different stress reactions compared to males (Young and Korszun, 2010). Indeed, one study has shown that classical conditioning using shock as the unconditioned stimulus, did increase neurogenesis in the ventral DG of females but not males (Dalla et al., 2009). Our results emphasize the importance of further investigation of sex differences in the preservation of neural stem cells in the hippocampus is a potential treatment (Briley et al., 2016).

### 4.3. The neural progenitor cell-type composition changes after mitosis with sex-dependent manner

Consistent with past studies, we found similar percentages of Sox2-ir cells and DCX-ir cells in the progenitor proliferating pool in male rodents (Sibbe et al., 2015; Nickell et al., 2017). However, we found that females had a greater increase in the percentages of BrdU-ir cells co-expressing DCX between two and 24 hours after mitosis whereas males did not exhibit any significant change between these two timepoints. This finding suggests that the neural progenitor cell-type composition within the actively dividing pool in females changes after each cell division more so than in males. It also suggests that early on in division, the daughter cells proceed more rapidly through the neuronal cell lineage in females compared to males. This finding may in part explain the ability of females to compensate for the lower levels of cell proliferation to end up with a similar number of new neurons at three weeks compared to males. More studies are needed to examine sex differences in the timeline and mechanism of the transition of proliferating progenitors to new neurons for a comprehensive understanding of the regulation of neural progenitor cell pool in males and females.

### 4.4. Neurogenesis in males has a different trajectory compared to females

The present study found that males, but not females, showed substantial changes in the density of BrdU-ir cells across timepoints with an early increase from 24 hours to one week followed by a substantial decrease from one to two weeks. The decrease was notable such that despite the fact that males showed greater density of one week old BrdU-ir cells than females, but there was no sex differences in density of older (two-three week) old BrdU-ir cells. Our findings are consistent with previous studies that demonstrating the same trajectory in male Sprague Dawley rats (Cameron et al., 1993; Snyder et al., 2009, 2012) and no significant sex difference in the amount of two week or three week old BrdU-ir cells in cage controls (Tanapat et al., 1999; Barha et al., 2011; Chow et al., 2013 but see Lee et al., 2014). Collectively these results suggest that males and females regulate adult neurogenesis differently as males produce more new cells and show greater attrition of these new cells, whereas females produce fewer new cells which are preserved across maturation. These findings may explain why spatial learning and or estrogens given during the first week of new neuron development increases the survival of new neurons in males, but not in females (Ormerod et al., 2004; Epp et al., 2007; Chow et al., 2013; Yagi et al., 2016). Taken together, these results suggest that spatial training between one week and two weeks after production of new neurons can prevent the attrition of adult-born neurons in males but perhaps not in females.

### 4.5. Males, compared to females, had greater cell proliferation in the dentate gyrus

Males had a greater density of Ki67-ir cells in the DG compared to females, consistent with findings in meadow voles (Galea and McEwen, 1999). In contrast a number of other studies have not found sex differences in cell proliferation in the DG (Lagace et al., 2007; Brummelte and Galea, 2010; Barha et al., 2011; Spritzer et al., 2017). However, these inconsistences may be related to estrous cycle, as only proestrous females show greater cell proliferation than male rats (Tanapat et al., 1999), although this effect has not always been noted (Lagace et al., 2007). None of the females in the Ki67 analysis were in proestrus and thus, we would expect lower levels of cell proliferation in these females. Consistent with our Ki67 results we also see increased BrdU-ir cells at 2h in males compared to females, but no differences at 24h, which likely has to do with the population that Ki67 labels versus the pulsatile BrdU (Kee et al., 2002).

### 4.6. Conclusion

In the present study, sex differences are noted in the neural stem cell population, cell proliferation, maturation rate and the attrition rate of adult-born neurons in the hippocampus. The trajectory of new neuron survival is dramatically different in males compared to females suggesting that the ability to influence neurogenesis within each sex may be due to the existing differences in timing and/or maturation of new neurons. Future studies should target mechanisms of these sex differences in adult neurogenesis as there are likely multiple factors involved that could profoundly affect these sex differences such as genetic (four core genotypes; 66), epigenetic (Sase et al., 2019) and mitochondrial functions (Biala et al., 2011) that differ between the sexes. These findings have profound implications for our understanding of adult neurogenesis in the DG, the use of therapeutics that modulate neurogenesis in the general population and underscore the need to include both sexes in research on hippocampal neurogenesis.

## Acknowledgements

We would like to thank Tanvi Puri, Yanhua Wen, and Stephanie Lieblich for the exceptional technical assistance with this work. This work was supported by a Natural Sciences and Engineering Research Council (NSERC) Discovery Grant to LAMG [RGPIN-2018-04301] and Izaak Walton Killam Memorial Pre-Doctoral Fellowship to SY.

## Notes

### Competing Interest Statement

The authors have declared no competing interest.

### Summary of Updates

New title, photomicrographs, some new analyses and data

